# Long chain monomethyl branched-chain fatty acid levels in human milk vary with gestational weight gain

**DOI:** 10.1101/2023.10.13.561203

**Authors:** Aifric O’Sullivan, Emer Brady, Lucy Lafferty, Fiona O’Shea, Zoe O’Regan, Noah Meurs, Michelle Baldini, Jivani Gengatharan, Christian M. Metallo, Martina Wallace

## Abstract

Breastfeeding is an important determinant of infant health and there is immense interest in understanding its metabolite composition so that key beneficial components can be identified. The aim of this research was to measure the fatty acid composition of human milk in an Irish cohort where we examined changes depending on lactation stage and gestational weight gain trajectory. Utilising a chromatography approach optimal for isomer separation, we identified 44 individual fatty acid species via GCMS and showed that monomethyl branched-chain fatty acids(mmBCFA’s), C15:0 and C16:1 are lower in women with excess gestational weight gain versus low gestational weight gain. To further explore the potential contribution of the activity of endogenous metabolic pathways to levels of these fatty acids in milk, we administered D_2_O to C57BL/6J dams fed a purified lard based high fat diet (HFD) or low-fat diet during gestation and quantified the total and *de novo* synthesized levels of fatty acids in their milk. We found that *de novo* synthesis over three days can account for between 10 and 50% of mmBCFAs in milk from dams on the low-fat diet dependent on the branched-chain fatty acid species. However, HFD fed mice had significantly decreased *de novo* synthesized fatty acids in milk resulting in lower total mmBCFAs and medium chain fatty acid levels. Overall, our findings highlight the diverse fatty acid composition of human milk and that human milk mmBCFA levels differ between gestational weight gain phenotypes. In addition, our data indicates that *de novo* synthesis contributes to mmBCFA levels in mice milk and thus may also be a contributory factor to mmBCFA levels in human milk. Given emerging data indicating mmBCFAs may be beneficial components of milk, this study contributes to our knowledge around the phenotypic factors that may impact their levels.

## Introduction

The importance of breastfeeding for both mother and infant health is well established [1] and there is intense interest in identifying the components of human milk and mechanisms that mediate this beneficial effect [2-4]. A primary question within this field is how genotype, phenotype and environment impact human milk composition and the consequences for infant health [5]. The impact of maternal obesity and excess gestational weight gain on milk composition and production is of particular relevance due to its rapidly increasing prevalence and its association with many adverse effects on offspring health such as increased risk for childhood obesity and metabolic syndrome as an adult [6]. Altered in utero development is an important mediator of this risk but experimental models have shown alterations in breastmilk composition can also contribute [7, 8], however detailed knowledge of how maternal obesity or gestational weight gain impacts human milk composition is lacking.

Human milk lipids account for nearly half of the total energy intake in young infants and human milk contains a diverse fatty acid composition that is important for many aspects of infant health such as neurodevelopment, immune function and cardiac maturation [9-11]. Dietary intake and endogenous metabolism are important drivers of milk fatty acid composition and mammary gland *de novo* lipogenesis is unique relative to other human lipogenic tissues due to the presence of type II thioesterase which facilitates *de novo* synthesis of large amounts of medium chain fatty acids [12]. Monomethyl branched chain fatty acids are also abundant in breastmilk [13, 14] and vary with geographical site [15, 16], lactation stage, gestational stage at birth [17] and are enriched in the sn-2 position of triacylglycerol in human milk [18]. These fatty acids have previously been shown to protect from necrotizing enterocolitis (NE) in rodent models of NE [19] and may be an important component of breastmilk that promotes infant health. Dietary consumption of mmBCFAs from food such as dairy is a significant source of mmBCFAs [15, 20], however previously we have shown that mmBCFAs can also be *de novo* synthesized in adipose depots via utilization of intermediates of branched chain amino acid metabolism by fatty acid synthase [21]. In addition, we have found that synthesis of these mmBCFAs in adipose depots is potently suppressed in genetically obese mouse models (ob/ob) and by a high fat diet. However, the contribution of *de novo* synthesis to mmBCFAs in milk and whether this is similarly impacted by increased weight gain is unknown.

The aim of this research was to measure the total fatty acid composition of human milk in an Irish cohort, examine changes depending on lactation stage and determine differences depending on gestational weight gain trajectory with a focus on mmBCFAs. In a follow-up study using a mouse model of increased gestational weight gain we examined the impact of diet on the amount of *de novo* synthesized mmBCFAs in milk.

## Materials and Methods

### Study design

The human data and samples reported in this paper were collected as part of the Healthy Start Study. The Research Ethics Committee at the National Maternity Hospital, Holles St, Dublin, and the University College Dublin Human Research Ethics Committee approved the study. All research was conducted in line with the principles outlined in the declaration of Helsinki, and written informed consent was obtained from all participants. The Healthy Start Study was a semi-longitudinal observation study that recruited healthy Caucasian pregnant mothers, living in Dublin, aged 18-45 years. Exclusion criteria included mothers with an underlying disease or those who experienced complications during pregnancy. All participants were attending antenatal maternity clinics in Holles Street Maternity Hospital between January and December 2014. Data were collected from mothers during pregnancy and at birth, milk and data was further collected at the end of month 1 and the end of month 2 as detailed below. Demographic and lifestyle data was recorded at the pregnancy study visit using a written questionnaire.

#### Anthropometric measurements and gestational weight gain

Mothers pre-pregnancy weight and height were self-reported. Weight and height were measured during pregnancy at the start of trimester 2 and the end of trimester 3 and again at the end of the month 2 postnatal time point. Weight was measured in duplicate using a Tanita body composition analyser BC - 420MA (Tanita Ltd, GB) after having voided, wearing light clothing and without shoes. Height was measured using the Leicester portable height measures (Chasmores Ltd, UK) with the participant’s head positioned in the Frankfurt Plane.

#### Gestational Weight Gain

Gestational weight gain (GWG) was calculated by subtracting measured weight at the start of trimester 2 from measured weight at the end of trimester 3. Adequacy of GWG was determined by comparing the actual GWG to the Institute of Medicine’s (IOM) 2009 GWG guidelines [22], using pre-pregnancy BMI to classify mothers. Recommended weight gain was calculated by multiplying the recommended rate per trimester by the number of weeks between recorded weights. Mothers were classified into 3 groups based on whether they were below, within or above the recommended GWG guidelines [22].

#### Human milk and formula sample collection

A 10-15 mL sample of breast milk or infant formula was collected at the end of month 1 and month 2 as previously described (Neville et. al., 1984). Breastfeeding mothers were asked to put their infant to one breast to allowing suckling for 2 min between 09:00 and 10:00 am. The mothers briefly stopped feeding and manually expressed 10-15 mL of breast milk into the collection bottle using their hands to stimulate and direct the flow of milk. The sample was placed on ice while normal feeding resumed. For infant formula collection, mothers were asked to prepare the infant formula as usual. Approximately 10-15 mL of the prepared formula was poured into the collection bottle. The date and time of collection, the brand (for formula samples), and the time since last feed was recorded by mothers. Samples were stored in the freezer and brought to the study centre on ice.

### Animal study

Animal handling and care followed the NIH Guide for Care and Use of Laboratory Animals. The experimental protocols were approved by the UCSD Institutional Animal Care and Use Committee. C57BL/6J mice were obtained from Jackson Laboratories. Age matched C57BL/6J male and female mice (which had given birth to one litter previously) were mated and then female mice were removed, singly caged and given either a lard based high fat diet (60% kcal fat, Envigo TD. 06414) or low fat diet (10% kcal fat, Envigo TD. 08806) throughout gestation and the first 10 days of lactation. All females and males were 15-17 weeks of age at the time of breeding. On day 7 post-parturition, dams were intraperitoneally injected with 0.035 mls/g body weight weight 0.9% NaCl-D_2_O and drinking water was replaced with 8% D_2_O enriched water. On day 10 post-parturition, pups were separated from dams 4 hours prior to breastmilk collection. Breastmilk was collected via manual expression of the milk following oxytocin (0.05 IU/g body weight, stock prepared as 10 IU/ml saline of Sigma O3251-500 IU) and ketamine/xylazine (100 mg/kg ketamine, 10 mg/kg xylazine) administration as previously described [23]. Following milk collection, blood was collected from the dam via decapitation and serum separated following clotting for 30 minutes by centrifugation @ 1500 g for 10 minutes at 4°C for D_2_O.

### GC/MS analysis of fatty acids

Total fatty acids were extracted from milk using a Folch-based methanol (MeOH) /chloroform (CHCl_3_)/saline extraction at a ratio of 1:2:1 with inclusion of [^2^H_31_] C16:0 as an internal standard for the murine milk and [^2^H_31_] C16:0, [^2^H_29_] C15:0 and [^2^H_3_] C18:0 used as internal standards for the human milk. Butylated hydroxytoluene was included in the CHCl_3_ at a concentration of 0.01% w/v. Briefly, 250 µl MeOH, 500 µls CHCl_3_, 250 µls saline and fatty acid isotope internal standards were added to 10 μls murine milk or 30 μls human milk. This was vortexed for 10 minutes followed by centrifugation at 10,000 g for 5 minutes at 4 °C. The lower CHCl_3_ phase was dried and then derivitised to form fatty acid methyl esters (FAMES) via addition of 500 µls 2% H_2_SO_4_ in MeOH and incubation at 50°C for 2 hours. FAMES were extracted via addition of 100 µl saturated salt solution and 500 µl hexane and these were analyzed using a Select FAME column (100m x 0.25mm i.d.) installed in an Aglient 7890A GC interfaced with an Agilent 5975C MS using the following temperature program: 80 °C initial, increase by 20 °C/min to 170 °C, increase by 1 °C/min to 204 °C, then 20 °C/min to 250 °C and hold for 10 min. The % isotopologue distribution of each fatty acid was determined and corrected for natural abundance using in-house algorithms adapted from Fernandez et al[24]. Fatty acids were quantified from deuterated internal standards or external standard curves and then expressed as mole % of total.

### Plasma and milk ^2^H_2_O enrichment analysis

The ^2^H labeling of water from samples or standards was determined via deuterium acetone exchange. 5 µl of sample or standard was reacted with 4 µl of 10N NaOH and 4 µl of a 5% (v/v) solution of acetone in acetonitrile for 24 hours. Acetone was extracted by the addition of 600 µl CHCl_3_ and 0.5 g Na_2_SO_4_ followed by vigorous mixing. 100 µl of the CHCl_3_ was then transferred to a GC/MS vial. Acetone was measured using an Agilent DB-35MS column (30 m 3 0.25 mm i.d. x 0.25 µm, Agilent J&W Scientific) installed in an Agilent 7890A gas chromatograph (GC) interfaced with an Agilent 5975C mass spectrometer (MS) with the following temperature program: 60 °C initial, increase by 20 °C/min to 100 °C, increase by 50 °C/min to 220 °C, and hold for 1 min. The split ratio was 40:1 with a helium flow of 1 ml/min. Acetone eluted at approximately 1.5 min. The mass spectrometer was operated in the electron impact mode (70 eV). The mass ions 58 and 59 were integrated and the % M1 (m/z 59) calculated. Known standards were used to generate a standard curve and plasma % enrichment was determined from this. All samples were analysed in triplicate.

To calculate the pup’s normalized % D_2_O enrichment as an indication of milk intake, the dam’s % D_2_O enrichment and the pup’s weight were taken into account using the following equation:

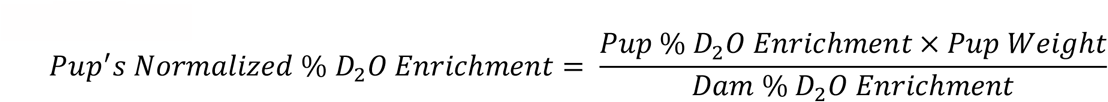

### *In vivo de novo* lipogenesis calculations

Calculation of the fraction of newly synthesized fatty acids (FNS) was based on the method described by Lee et al. [25], where FNS is described by the following equation:

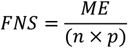

Where ME is the average number of deuterium atoms incorporated per molecule (ME =1 x *m*1 + 2 x *m*2 +3 x *m*3 …), p is the deuterium enrichment in the dams plasma and n is the maximum number of hydrogen atoms from water incorporated per molecule. The value for n was based on previous publications indicating 22 deuteriums are incorporated into palmitate *in vivo* in rodents using a similar dosing protocol as this study [25]. Based on this it was reasoned that 3 deuterium could be incorporated per 2 carbon elongation due to direct incorporation from water, through NADPH or from acetyl-CoA and 1 deuterium could be incorporated into the initial acyl-ACP from acetyl-CoA. For fatty acids whose synthesis is not initiated via a 2 carbon acyl unit derived from acetyl-CoA but rather from propionyl-CoA (odd chain fatty acids) or branched-CoAs (mmBCFAs), it was assumed that there was no deuterium present in the initial acyl-ACP. The n number used for each fatty acid is shown in Supplementary table 1.

| Fatty acid | n |
| --- | --- |
| C12:0 | 16 |
| C14:0 | 19 |
| C14:1 | 17 |
| iso C15:0 | 12 |
| anteiso C15:0 | 12 |
| C15:0 | 18 |
| iso C16:0 | 18 |
| C16:0 | 22 |
| C16:1 | 20 |
| iso C17:0 | 18 |
| anteiso C17:0 | 18 |
| C17:0 | 21 |

The molar amount of newly synthesized fatty acids (MNS) was determined by:

MNS = FNS x total fatty acid amount (nM).

### Statistics

Normal distribution of data was tested using the Shapiro-Wilk test in SPSS. Differences in fatty acid levels were determined via a two tailed t-test with a false discovery rate of 5% determined via the Two-stage step-up (Benjamini, Krieger, and Yekutieli) method in Prism 9 (La Jolla, CA). In addition, a general linear model including age as a covariate was carried out in SPSS. Significant differences in animal studies was calculated using a two tailed t-test and significance was determined as p<0.05 (Prism 9, La Jolla, CA).

## Results

### Participant characteristics

In total, 62 mothers were recruited to the study. One mother dropped out before giving birth, 8 dropped out before the end of month 1 and 5 more dropped out before the end of month 2, leaving a cohort of 48 mothers for this analysis. All mothers were Caucasian, aged between 25 and 42 years (mean age, 33 ±3.6 years) with BMI in the normal range prior to pregnancy (22.9 ±2.5 kg/m2). Most mothers finished third level education (80%), were married (87%) and employed (85%). Almost half of the participants were first time mothers (48%), the majority had a vaginal delivery (83%) at an average of 40.3 ±1.2 weeks gestation. On average mothers gained 11.3kg ±3.9 kg during their pregnancy. However, when weight gain guidelines were individualized based on the Institute of Medicine (IOM) criteria which takes into account pre-pregnancy BMI, only 40% stayed within their target range. Seventeen percent of women gained less weight than recommended and 44% exceeded weight gain recommendations during their pregnancy.

### mmBCFAs decrease with increasing gestational weight gain in human milk

Of the 48 mother-infant pairs included in this analysis, 37 mothers providing a breastmilk sample at 1 month post parturition, and 31 of these provided a second sample at month 2. 11 mothers were formula feeding at month 1 post-parturition and provided formula samples. The demographics and phenotypic characteristics of mothers who provided breastmilk samples are presented in Table 1. GWG values were missing for 4 mothers and of the remaining 33 samples, 46% (n=15) stayed within their targeted GWG, 21% (n=7) were below and 33% (11) were above. Using an untargeted GCMS approach, 44 fatty acids were detected in human milk samples taken at 1 month and 2 months post parturition (table 2). This included 6 monomethyl branched-chain fatty acid species that account for an average of 0.5% of fatty acids in breastmilk (table 2). Paired analysis of changes in the mole % composition of fatty acids in the breastmilk between month 1 and month 2 indicated a significant decrease in C12:0, C14:0, C22:2 and total medium chain fatty acids (table 2). As 11 of the originally recruited mothers were primarily feeding formula by the end of month 1, we also collected formula samples which spanned 6 different brands available on the Irish market at that time and compared this with the human milk fatty acid composition. Variation in fatty acid levels in the formula samples was less than that of the human milk samples (Table 2). Comparison of the mole % fatty acid composition of all formula samples with breastmilk collected in month 1 indicated extensive differences in fatty acid composition with some species undetectable in formula (C14:1, C20:3n3, C22:2). The biggest fold change decrease in formula versus breastmilk was found in mmBCFAs, C20:3n6, C16:1n10(sapienate) and C16:1n7 and the biggest fold change increase was found in the saturated fatty acids C20:0 – C24:0 (Fig 1b). However, there was no significant difference in total monounsaturated fatty acids, polyunsaturated fatty acids or the essential fatty acids EPA (C20:5n3) and DHA (C22:6n3) (Table 2).

**Table 1.** Demographics of mothers who donated breastmilk (n=37).

|  | Mean (SD) |
| --- | --- |
| Age (years) | 33 (3.6) |
| Trimester 1 weight (kg) | 61.9 (8.0) |
| Trimester 1 BMI (kg/m<sup>2</sup>) | 22.7 (2.6) |
| Trimester 3 weight (kg) | 73.1 (9.1) |
| GWG (kg) | 11.3 (3.9) |
| Below GWG recommendations | 21.2 % |
| Within GWG recommendations | 45.5 % |
| Above GWG recommendations | 33.3 % |
| Gestational weeks | 40.5 (1.2) |
| Infant gender (% Female) | 54 % |

GWG recommendations refer to the Institute of Medicine's recommendations for gestational weight gain, based on pre-pregnancy BMI[22].

**Table 2.**
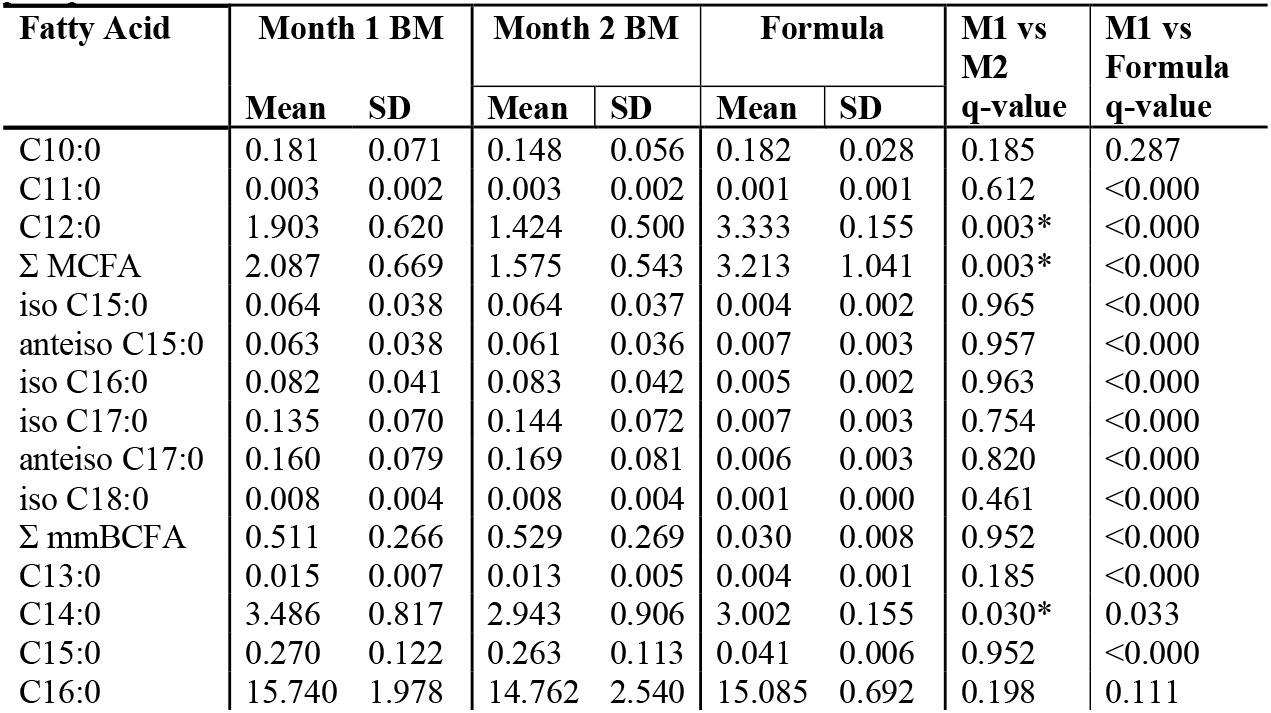
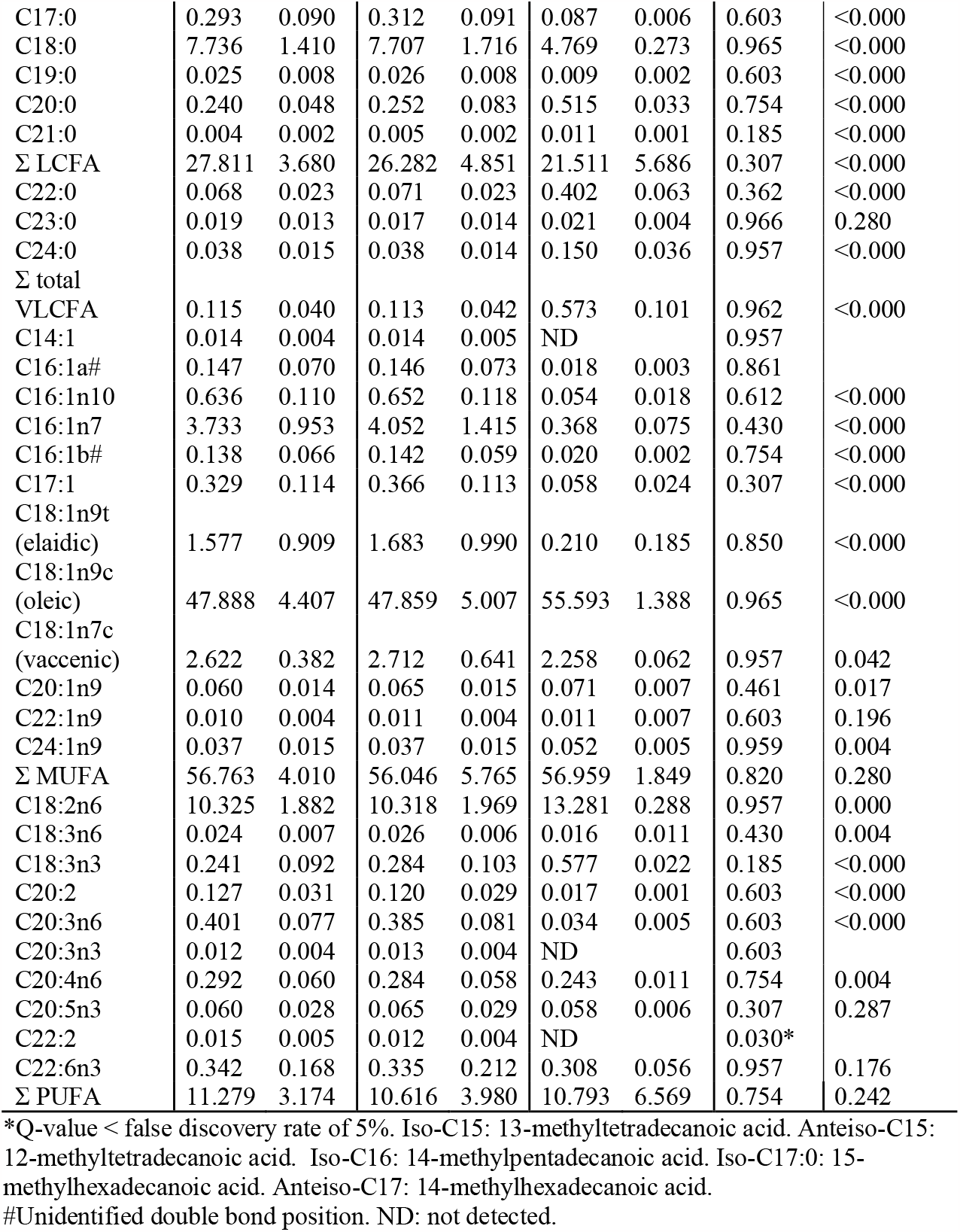
Mol % Fatty acid composition of breastmilk samples taken 1 month or 2 months post parturition.

**Figure 1.**
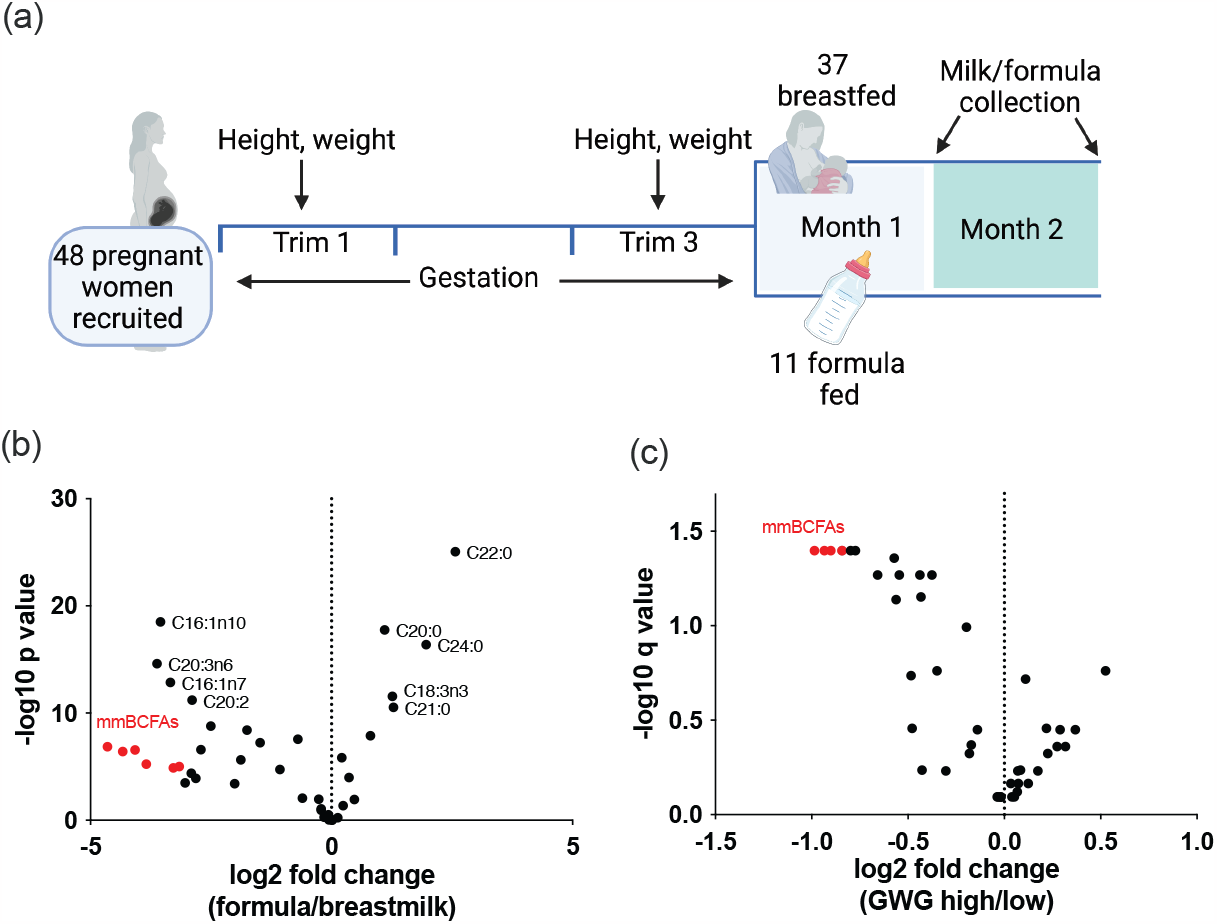
mmBCFAs decrease with increasing gestational weight gain in human milk. a) Overview of human milk collection study design. Volcano plot showing the difference in fatty acid composition between b) formula and human milk c) milk from women with high or low gestational weight gain.

To determine how weight gain during gestation impacts mmBCFAs in breastmilk, we compared samples from women who gained above (high GWG), below (low GWG) or within the recommended gestational weight gain as recommended by the institute of medicine [22]. We found that when we compared samples from those with high GWG to those with low GWG, mmBCFAs, C15:0 and a C16:1 species were the most significantly decreased fatty acids in the high GWG group (Fig 1c and d, Table 3). These data indicate that levels of BCFAs in breastmilk are altered with gestational weight gain and that excessive weight gain may modulate BCFA levels in breastmilk in a similar way as noted in response to obesity [26-28]. However, there were minimal significant differences when comparing across all three GWG groups, or when low GWG or high GWG was compared with breastmilk from those who gained within the recommended GWG. This may be due to the relatively small size of the cohort when divided by GWG and the phenotypic range (Table 4).

**Table 3.**
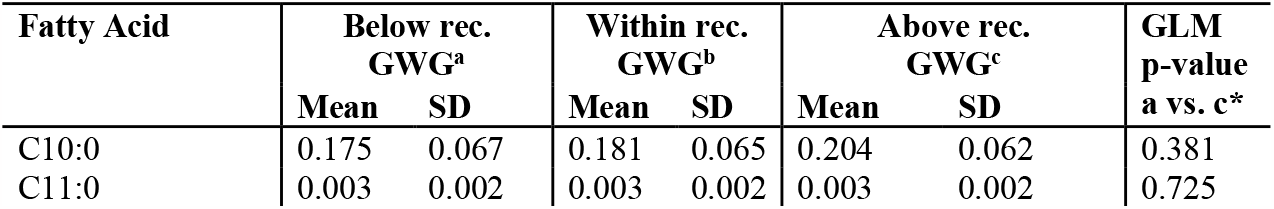
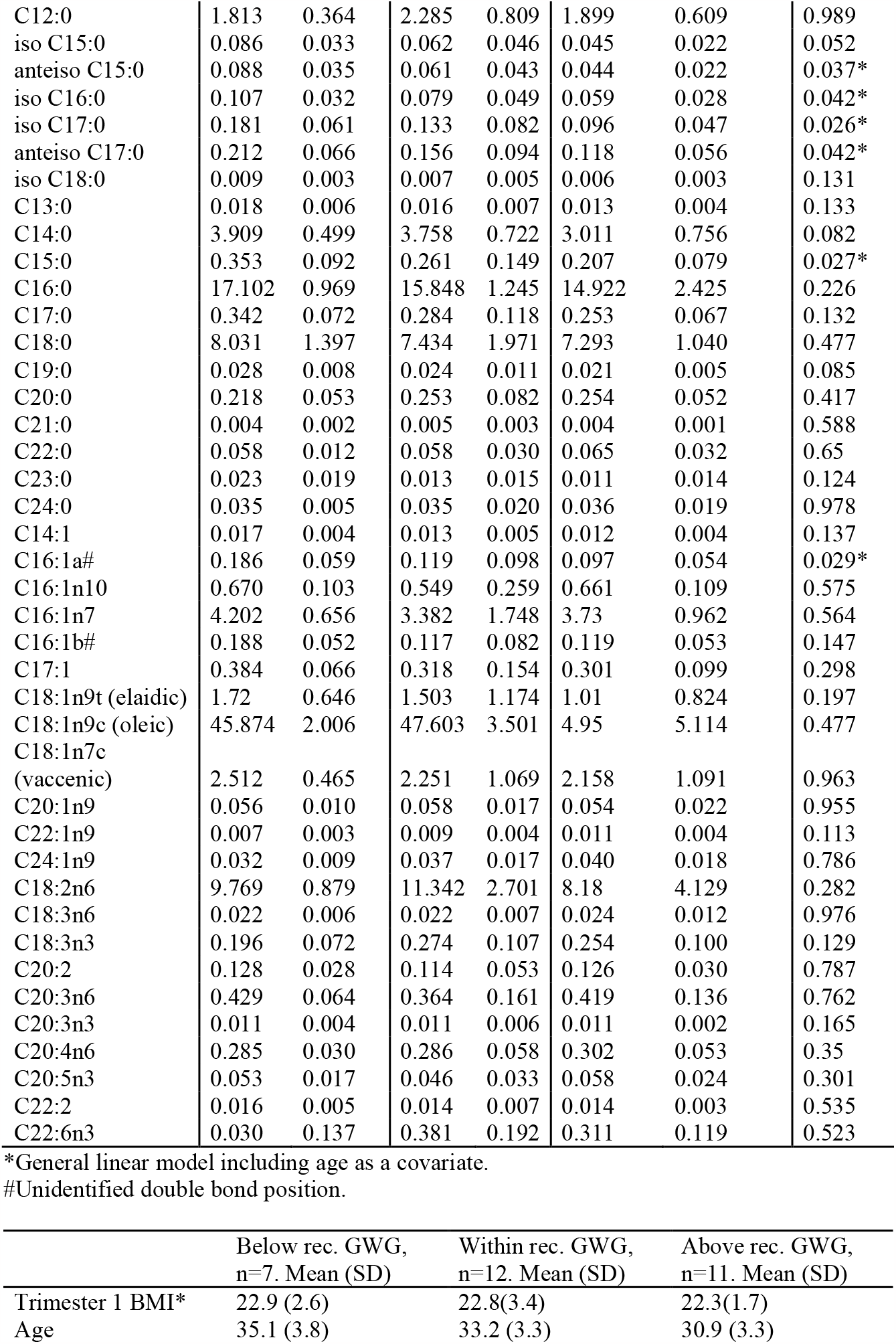
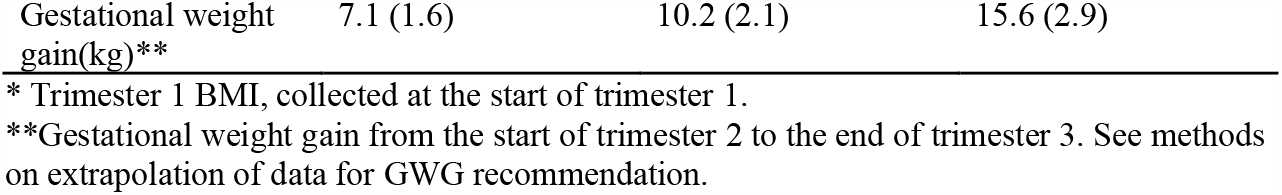
Mol % Fatty acid composition of breastmilk samples taken from women who gained above or below gestational weight gain recommendations.

### *De novo* synthesized BCFAs in murine milk are decreased by a high fat diet

mmBCFA levels in human milk vary with dietary intake of food containing mmBCFAs indicating dietary intake is an important determinant of their levels [15]. However, our previous studies have indicated that *de novo* synthesis from branched chain amino acids can also contribute to mmBCFA levels in humans and that mmBCFA synthesis is sensitive to obesity induced changes in the total activity of *de novo* lipogenesis [21]. To establish whether *de novo* synthesis is a contributor to mmBCFA levels in mammalian milk and thus may also contribute to human milk levels, we administered D_2_O to C57BL/6J dams from days 7-10 post-parturition and quantified the *de novo* synthesis rate of a range of fatty acids present in their milk. In addition to this, we pre-fed these dams during gestation and lactation with either a high fat (HFD) or low fat diet (LFD) to determine whether known dietary modulators of *de novo* lipogenesis impact synthesis of mmBCFAs (Fig 2a). Only 43% of LFD mice and 40% of the HFD mice had litters following the switch from a chow to a defined diet. Of these litters, 50% of control and 56% of HFD litters survived to P10 (Fig 2b). This is in contrast to these dams on their first breeding when maintained on normal vivarium chow where 93% had litters and 50% of these litters survived to day 10. These initial results indicate that changing the diet to a defined diet, whether high fat or low fat during the beginning of gestation is detrimental to breeding success. HFD fed dams were significantly heavier on day 0/1 post parturition(Fig 2c) and there was no significant differences in litter size (Fig 2d). However, milk intake was decreased in the pups of dams fed a HFD as indicated by deuterium body water enrichment in the pups[29](Fig 2e).

**Figure 2.**
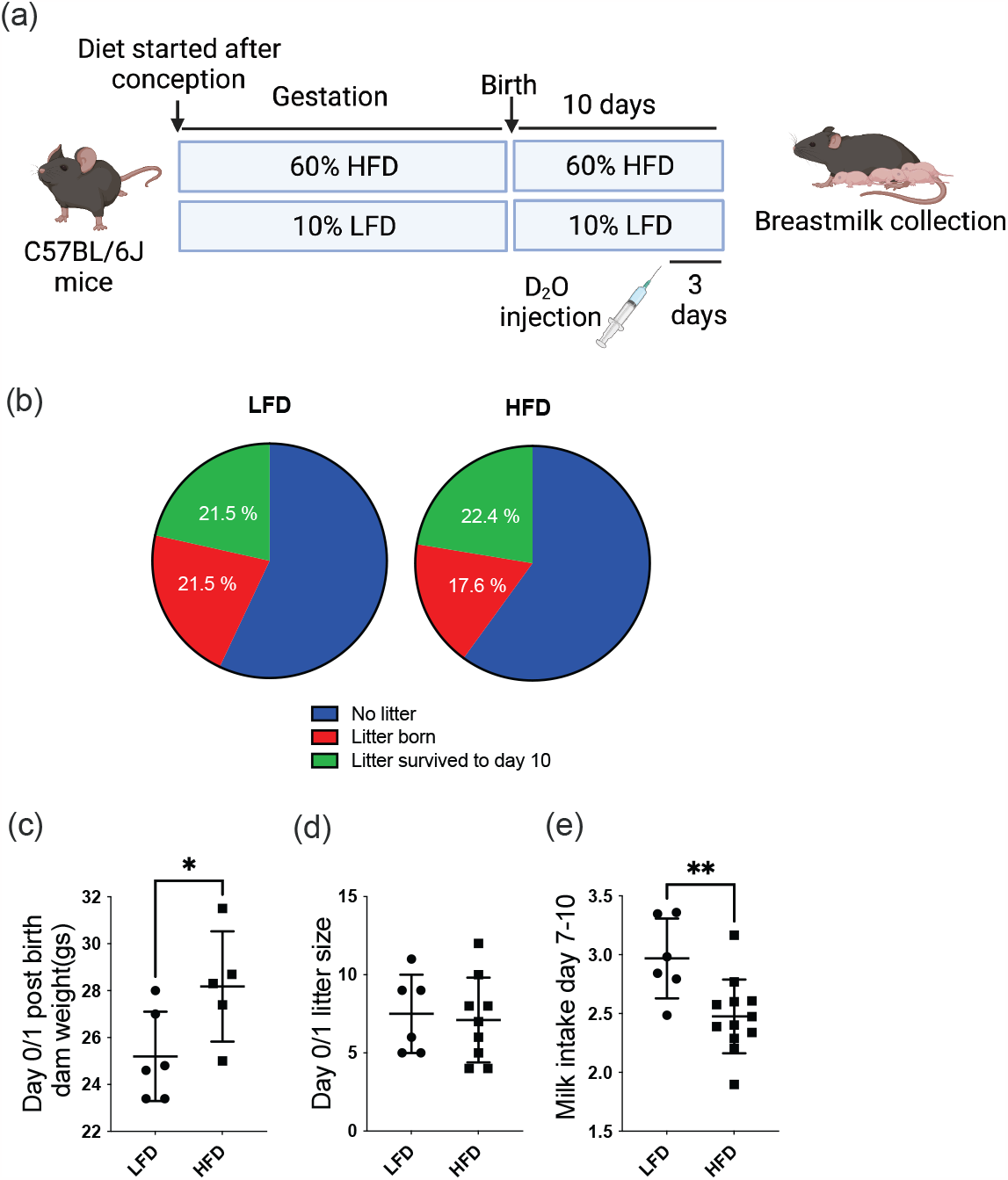
The impact of feeding a high fat diet (HFD) or low fat diet (LFD) during gestation on dam weight and breeding success. A) overview of study design b) the % of litters born and survival to day 10 post birth C) Dam weight following birth (LFD n=6, HFD n=5) d) litter size e) pup milk intake determined by normalized body water enrichment from deuterium. ^*^=p<0.05, ^**^=p<0.01, ^***^ =p<0.001.

We next determined whether the fatty acid composition of the milk changed with diet. HFD fed dams had lower amounts of fatty acids with chain lengths of 12-16 but higher amounts with chain lengths of C17-C20 (Fig 3a-b). Levels of mmBCFAs also followed this trend, with shorter chain BCFAs (C15 – C16) decreasing and longer chain BCFAs (C17) increasing with HFD. However when we quantified the amount of fatty acids that were *de novo* synthesised, we found a general decrease with high fat diet across across all fatty acids (Fig 3c-f). Next we focused on the fractional contribution of *de novo* synthesis to each fatty acid and found that *de novo* synthesis contributed in the range of 10-50% to mmBCFA species in dams who were fed the low fat diet with higher synthesis levels in the shorter mmBCFA species (Fig 3g). When combined, the data indicates that levels of fatty acids with chain lengths of 12-16 are likely more driven by *de novo* lipogenesis and longer chain fatty acids are being supplied by the diet or fatty acids synthesized and stored prior to D_2_O administration and/or gestation. Of note, previously we quantified iso-C16, iso-C17 and anteiso-C17 in the HFD utilised here and found levels of mmBCFAs with a chain length of C17 higher than that of iso-C16, thus HFD fed mice will consume more of these longer chain mmBCFAs[21].

**Figure 3.**
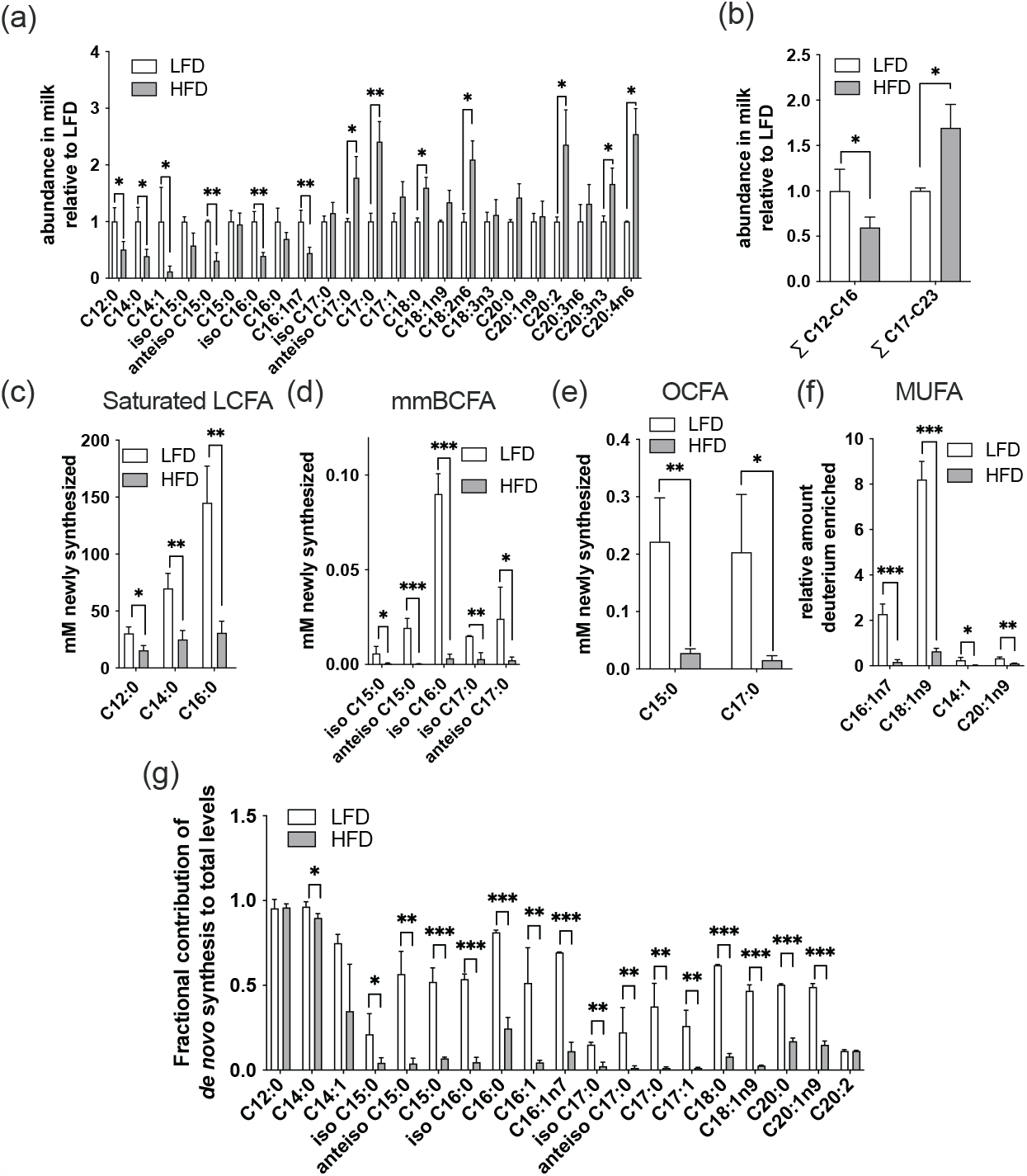
De novo synthesized BCFAs are decreased by high fat diet in murine milk. a-b) relative fatty acid levels in milk taken at day 10 post birth from HFD versus LFD fed mice. c-f) levels of de novo synthesized fatty acids in milk from HFD versus LFD fed mice. (LFD n=2, HFD n=4) ^*^=p<0.05, ^**^=p<0.01, ^***^ =p<0.001.

Overall, our findings indicate that levels of *de novo* made fatty acids in murine milk is suppressed by a high fat diet, similarly to what happens in adipose tissue in response to a high fat diet [21]. Due to the unique metabolic adaptions of fatty acid synthesis in the mammary gland that generates both medium and long chain fatty acids, this results in a decrease in fatty acids with chain length C12 – C16. This data highlights how metabolic adaptions to dietary composition and obesity can influence breastmilk composition via alterations in *de novo* lipogenesis.

## Discussion

Maternal obesity and gestational weight gain are important determinants of offspring health [30]. Given that obesity is now commonly recognized as an ongoing global health pandemic [31] that results not just in diseases associated with the metabolic syndrome but also increases risk of mortality from infectious diseases and cancer [32, 33], there is an increasing need to understand how altered weight gain impacts health during all parts of the life cycle. Here, we set out to determine how altered weight gain during gestation impacted the fatty acid composition of breastmilk as obesity induced alterations in breastmilk composition are known to impact offspring health.

We found that mmBCFAs were the most decreased fatty acids in women who gained above the recommended gestational weight gain compared with those that gained below. In addition, we found that in a mouse model of increased gestational weight gain, synthesis of mmBCFAs was significantly decreased and levels of mmBCFAs with a chain length < C17 were also decreased. Previously we have found that a significant fraction of plasma mmBCFAs in humans is *de novo* synthesized and we and others have found that mmBCFAs are decreased with obesity in both humans and obese mouse models and correlate with insulin sensitivity[21, 26-28]. Collectively this data indicates that mmBCFA levels may respond similarly in breastmilk and synthesis may decrease with increasing weight gain leading to decreased levels in breastmilk. As previous studies have shown mmBCFAs are protective in a rodent model of necrotizing enterocolitis and impact immune cell function and microbiota composition[19], exposure to increased mmBCFAs via breastmilk may be advantageous to offspring health. In addition, recent studies have shown that mmBCFA containing lipids from microbiota can modify host immune responses differentially compared to lipid ligands containing other acyl chains further indicating mmBCFAs may impact infant immune development [34, 35]. However, further studies will be required to determine if enhancing mmBCFA levels in breastmilk in the context of obesity may attenuate some of the detrimental impacts on offspring health.

This study also highlighted the impact of a high fat diet on the amount of *de novo* synthesized fatty acids present in milk. In our mouse model, we found a high fat diet resulted in a general decrease in saturated, monounsaturated, branched and odd chain fatty acids. This translated to a decrease in levels of fatty acids primarily with a chain length of <17 highlighting the importance of *de novo* synthesis to medium chain fatty acid levels in breastmilk, whereas longer chain fatty acid levels are likely to be driven more by diet or endogenous fatty acids stored in adipose tissue. Previous studies in humans with ^13^C glucose also indicated that de novo synthesis primarily contributed to synthesis of fatty acids with a chain length <C17 in human milk and that high fat feeding decreased *de novo* synthesis [36]. Notably, we did not find a change in medium chain fatty acids in human milk in the increased gestational weight gain group which may be due to the limited size and phenotypic range of our cohort or that synthesis of these fatty acids may be more related to the macronutrient composition of the diet rather than weight gain.

Finally, our analysis also highlighted the diversity of fatty acids species in breastmilk with 44 individual fatty acid species reported. The fatty acid composition of breastmilk reflects dietary intake, endogenous synthesis by the mammary gland and fatty acids released from other tissue depots [37]. Considering that many of these fatty acids are not typically reported in plasma, it is likely that local metabolism in the mammary gland is an important factor in determining this diversity. Indeed, the presence of the enzyme TE2 in mammary glands ensures production of medium chain fatty acids as it interacts with FASN to cause release of shorter chain fatty acids than its typical product palmitate[12]. Another notable characteristic of breastmilk is the diversity of monounsaturated fatty acids with 4 distinct monounsaturated C16 fatty acids identified. Desaturation of fatty acids is primarily carried out by the enzymes SCD1, FADS1 and FADS2. FADS2 demonstrates the most tissue specificity with expression highest in the skin where it is responsible for production of sapienic acid and monounsaturated odd chain fatty acids which are also found in breastmilk[38]. In line with local production playing an important role, FADS2 is the only desaturase upregulated with days postpartum in mRNA from human milk fat globule[39]. Overall, this data shows that breastmilk has a significantly diverse range of fatty acids which is likely influenced by both endogenous metabolism and diet.

In conclusion, this study is the first to our knowledge that determined how mmBCFA levels in breastmilk alter with gestational weight gain and to quantify the amount of *de novo* made mmBCFAs in mammalian breastmilk. This is an important step in further understanding the role of mmBCFAs in mammalian physiology and how excess gestational weight gain impacts milk fatty acid composition which may have longer term impacts on infant health. However, there are a number of limitations with this study as it was primarily focused on gestational weight gain and not analysis of milk from mothers with pre-gestational obesity which may have a more pronounced impact. In addition, dietary intake is also an important driver of mmBCFA levels and increased dietary dairy fat has been shown to increase mmBCFA levels in human milk[40]. To further understand the association between weight gain and mmBCFA levels in human milk, further studies in larger cohorts of women with pre-gestational obesity and collection of dietary intake data would be required. This will lay the groundwork for determining whether mmBCFA levels in breastmilk are a modifiable factor that could have a beneficial impact on infant health.

## Author contributions

A.O’S conceived and designed human milk study, C.M.M and M.W. conceived and designed animal studies. A.O’S, Z.O’R, and F.O’S collected human milk samples. M.W, N.M, J.G, A.O’S, E.B, L.L, Z.O’R and F.O’S analysed human milk samples, M.W. carried out animal experiments and M.W. and M.B. analysed animal samples. A.O’S and M.W wrote the manuscript with help from all authors.

## Acknowledgements and Funding

This work was supported by a Seed grant made available through the UC San Diego Larsson-Rosenquist Foundation Mother-Milk-Infant Center of Research Excellence. to M.W and through the UCD Seed Funding .Scheme to A.O’S. The support of the Family Larsson-Rosenquist Foundation is gratefully acknowledged.

## Data availability

The data that support the findings of this study are available from the corresponding author M.W., upon reasonable request.

